# Does Pre-Sensorimotor Brain Activity Modulate TMS-Evoked Potentials?

**DOI:** 10.64898/2026.09.10.750602

**Authors:** Motahare Delbari, Christoph S. Herrmann, Christian Grefkes, Stefan Debener

## Abstract

Transcranial magnetic stimulation combined with electroencephalography (TMS–EEG) provides a direct measure of cortical responses to local brain perturbations and offers a powerful approach for investigating brain-state-dependent neural dynamics. However, it remains unclear to what extent ongoing sensorimotor brain activity influences TMS-evoked potentials (TEPs).

To address this question, thirty healthy participants performed kinesthetic motor imagery of left- and right-index finger tapping while single-pulse TMS was applied over the left primary motor cortex. 64-channel EEG was recorded to characterize sensorimotor oscillatory activity and TEPs. Time–frequency analyses quantified event-related desynchronization (ERD) in the mu (8–12 Hz) and beta (13–30 Hz) frequency bands, while TEP amplitudes were examined within early latency windows (10-65 ms).

Motor imagery induced robust bilateral mu- and beta-band ERD, confirming successful engagement of the sensorimotor network. However, ERD patterns were highly similar between left- and right-hand motor imagery, and neither frequentist nor bayesian analyses identified differences in TMS-evoked potentials between imagery conditions. Source reconstruction demonstrated rapid propagation of TMS-evoked activity from the stimulated motor cortex to distributed bilateral motor regions. Correlation analyses revealed that greater pre-stimulus beta-band ERD was associated with larger P60 amplitudes. Because the instructed motor imagery conditions did not produce distinguishable pre-stimulus brain states, an exploratory trial-by-trial analysis was subsequently performed to account for naturally occurring variability in sensorimotor lateralization to investigate the effect of variability in ERD on TRP. This analysis revealed an overall effect of trial-by-trial brain state on TEP amplitudes.

## Introduction

Transcranial magnetic stimulation (TMS) enables the non-invasive induction of neural activity in the human brain and has become a well-established method for probing cortical excitability and connectivity (Barker et al., 1985; Di Lazzaro & Rothwell, 2014). Single TMS pulses applied over the primary motor cortex (M1) can elicit motor-evoked potentials (MEPs), which are thought to reflect corticospinal excitability; at the peripheral level electromyography (EMG) responses of target muscles can be observed. The combination of TMS with concurrent electroencephalography (EEG) complements this approach by directly capturing cortical responses to stimulation. TMS-evoked potentials (TEPs) reflect the spatiotemporal activation of distributed neural circuits and provide electrophysiological markers of cortical reactivity, inhibition, and effective connectivity (Ilmoniemi et al., 1997; Miniussi & Thut, 2010). TEPs are increasingly used to assess motor network function in both healthy individuals and clinical populations (Biabani et al., 2021; Rogasch & Fitzgerald, 2013; Tscherpel et al., 2020). In stroke patients, alterations in specific TEP components have been associated with motor network disruption and recovery potential (Sarasso et al., 2020; Tscherpel et al., 2020). These studies highlight the value of TEPs as a biomarker of cortical integrity and functional reorganization.

Despite its growing clinical and scientific application, TEPs exhibit substantial trial-to-trial as well as intra- and inter-individual variability. A significant proportion of this variability may be related to fluctuations in ongoing neural activity rather than variability in stimulation parameters alone. Oscillatory brain states have emerged as an important factor influencing cortical responsiveness, highlighting the need to consider the state of the brain at the time of stimulation when interpreting TMS-evoked responses (Arieli et al., 1996; López-Alonso et al., 2014). Brain-state dependence has been studied in MEPs using EEG-triggered TMS (Bergmann, 2018; Desideri et al., 2019; Hussain et al., 2019; Karabanov et al., 2021; Poorganji et al., 2023; Sack et al., 2023; Takemi et al., 2013; Zrenner et al., 2015). These studies have demonstrated that corticospinal responses vary as a function of the phase and power of ongoing sensorimotor oscillations, with stimulation delivered during specific phases of the mu-rhythm producing systematically different MEP amplitudes. However, whether similar state-dependent effects extend to cortical responses like TEPs remains less clear. Although MEPs and TEPs are both elicited by the same TMS pulse, they reflect distinct physiological processes (Siebner et al., 2022). While MEPs represent the readout of the corticospinal system, TEPs are thought to reflect the activation and propagation of activity within distributed cortical networks. Consequently, brain states – as reflected by oscillatory brain activity – may influence corticospinal and cortical responses differently. To investigate whether mentally induced changes in sensorimotor state influence cortical TMS responses, a procedure that modulates sensorimotor oscillatory activity without introducing movement artifacts is needed.

The present study combined a mental task, motor imagery (MI), with concurrent TMS–EEG to investigate whether sensorimotor brain state fluctuations influence cortical responses to TMS. Kinesthetic MI engages sensorimotor networks and induces event-related desynchronization (ERD) in the mu (8–12 Hz) and beta (13–30 Hz) frequency bands over sensorimotor cortex (Pfurtscheller & Lopes Da Silva, 1999; Pfurtscheller & Neuper, 1997). ERD induced by MI is widely interpreted as reflecting reduced neuronal synchrony and increased cortical activation (Jeon et al., 2011; Pfurtscheller, 2001). While MI produces robust ERD effects and is frequently used for neurofeedback and brain– computer interface applications (Zich, Debener, et al., 2015, 2017; Zich, De Vos, et al., 2015; Zich, Harty, et al., 2017), the strength of sensorimotor desynchronization is known to vary both within and across participants (Holper et al., 2012). We therefore used MI as an experimental task to examine whether fluctuations in sensorimotor oscillatory activity are associated with variability in TMS-evoked potentials (TEPs). To induce variability in sensorimotor brain state, participants performed left- and right-hand motor imagery, and we tested whether these conditions differentially modulated TEP amplitudes. This was done for early M1-TEP responses, which are known to indicate different motor network processing states (Hernandez-Pavon et al., 2023; Ilmoniemi et al., 1997; Ilmoniemi & Kičić, 2010; Rogasch & Fitzgerald, 2013). We further hypothesized that pre-stimulus mu- and beta-band activity over the stimulated primary motor cortex would be associated with variability in TEP components, particularly early responses reflecting motor cortical processing. Finally, because motor imagery does not consistently produce strongly lateralized sensorimotor activation (Zich, Harty, et al., 2017), we conducted an exploratory analysis in which trials were classified according to a sensorimotor laterality index derived from mu–beta ERD. This trial-by-trial approach was intended to determine whether accounting for endogenous fluctuations in brain state could reveal modulation of TMS-evoked responses not captured by the instructed imagery conditions.

## Methods

### Participants

Thirty healthy adults participated in this study (24 female, age: 18-30 years, mean age 24.8 ±2.9 years), all right-handed with normal or corrected vision. Two datasets were excluded due to technical issues with the EEG recording, resulting in a final sample of twenty-eight participants. Participants were screened for TMS contraindications. Exclusion criteria included pregnancy, metallic implants in the head, cardiac pacemakers, epilepsy, and neurological or psychiatric disorders affecting the central nervous system. Handedness was confirmed using the Edinburgh Handedness Inventory (Oldfield, 1971). Motor imagery ability was assessed using the short version of the Kinesthetic and Visual Imagery Questionnaire (Malouin et al., 2007). Depressive symptoms were screened using the Beck Depression Inventory-II (BDI-II), and general health status was assessed using the Short Form Health Survey (SF-12). The study design, hypotheses, and primary analyses were preregistered prior to data analysis (OSF: https://doi.org/10.17605/OSF.IO/2HD6Y). Any deviations from the preregistered analysis plan will be reported.

### TMS-EEG Recording

TMS was delivered using a PowerMAG Research 30 stimulator (MAG & More GmbH, Munich, Germany) equipped with a 70-mm figure-of-eight coil generating biphasic pulses. The coil was positioned over the left primary motor cortex (M1), corresponding to the cortical representation of the right index finger. Coil position was continuously monitored using Visor2™ neuronavigation software to ensure accurate targeting throughout the experiment. EEG was recorded using a TMS-compatible 64-channel system (BrainAmp DC, Brain Products GmbH, Germany). Ag/AgCl sintered ring electrodes were mounted in an elastic cap (EasyCap GmbH, Germany) using an equidistant electrode montage. Electrode impedances were kept below 5 kΩ. The reference electrode was positioned at the tip of the nose, and the ground electrode between AFz and Fz. Vertical and horizontal eye movements were monitored using electrodes placed below eyes. EEG signals were continuously recorded at a sampling rate of 5 kHz with a resolution of 0.1 μV/bit and were online filtered between 0.1 and 1 kHz. EEG was recorded continuously during TMS using a TMS-compatible amplifier that prevented saturation of the EEG signal following stimulation.

### Experimental Procedure

Participants were seated comfortably in an armchair in a quiet laboratory environment. A computer monitor was positioned approximately 1.2 m in front of the participant at eye level, and both hands rested comfortably on the armrests. Prior to the main experiment, participants completed 10 practice trials to familiarize themselves with the task and experimental procedure. The session began with a 10 minutes resting-state EEG recording during which participants were instructed to maintain visual fixation on a cross presented on center of the monitor with their eyes open. Resting motor threshold (RMT) was then determined over the left primary motor cortex (M1). Single TMS pulses were delivered beginning at 40% of maximum stimulator output (MSO) and gradually increased until motor-evoked potentials (MEPs) with a peak-to-peak amplitude of at least 50 μV were elicited in at least 5 out of 10 consecutive trials. The optimal stimulation site (motor hotspot) was defined as the scalp location that consistently produced the largest MEPs in the right first dorsal interosseous (FDI) muscle at the lowest stimulation intensity. Following threshold determination, single-pulse TMS was delivered at 80% of the individual RMT (subthreshold intensity), and no overt motor responses were observed during the experiment. The TMS coil was positioned tangentially to the scalp at approximately 45° backwards relative to the midline. A coil spacer was placed between the coil and scalp to reduce mechanical vibration artifacts and minimize TMS-related electrode movement, thereby improving EEG signal quality (Ruddy et al., 2018). EMG activity was continuously recorded from the right FDI muscle throughout the experiment.

Participants then performed two kinesthetic motor imagery (MI) conditions: imagined right-index finger tapping and left-index finger tapping. The imagery conditions were presented in a fixed ABBA block sequence (A = right-hand MI, B = left-hand MI), which was identical for all participants. Each trial began with a fixation period of 3–5 s (uniformly jittered), followed by a visual cue presented for 1 s. The cue indicated the upcoming imagery condition and provided a brief preparation period before imagery onset. When the letter “R” appeared on the screen, participants were instructed to kinesthetically imagine performing repetitive tapping movements with their right index finger. Likewise, when the letter “L” appeared, they were instructed to kinesthetically imagine performing repetitive tapping movements with their left index finger. Participants were asked to maintain the imagined movement continuously for 5 s without producing any overt muscle contractions. A single TMS pulse was delivered at a random time point between 2 and 3.5 s after imagery onset. Participants were instructed to continue the imagery task despite the TMS pulse and to remain focused on the imagined movement until the end of the trial (5 s). The experiment consisted of 20 blocks of 10 trials each, resulting in 100 trials per imagery condition (200 trials in total) and 10 s rest period was provided between blocks. To reduce auditory contamination of the EEG recordings, participants wore earplugs throughout the experiment. The experimental session lasted approximately 40 minutes.

### Data Analysis

EEG preprocessing was performed in MATLAB R2022b (The MathWorks, Natick, MA, USA) using EEGLAB (Delorme & Makeig, 2004) and the TESA toolbox (Rogasch et al., 2017). Noisy channels were identified using a kurtosis-based criterion (5 standard deviations) and removed. Data were segmented into epochs ranging from −5000 to 2000 ms relative to the TMS pulse and baseline corrected using the interval from −5000 to −4000 ms (Rogasch et al., 2017). The TMS pulse artifact (−2 to +10 ms) was removed and reconstructed using cubic interpolation. Data were subsequently downsampled to 1000 Hz. Trials contaminated by large-amplitude artifacts were identified and rejected using a joint probability criterion (5 standard deviations). EEG data were high-pass at 1 Hz and low-pass filtered at 80 Hz using a fourth-order Butterworth filter. A notch filter (48–52 Hz) was applied to attenuate line noise. To attenuate residual TMS-related, ocular, and muscle artifacts, Fast Independent Component Analysis (FastICA) was performed. Artifact-related components were identified using the automated component classification procedures implemented in TESA and subsequently verified by visual inspection before removal (mean ± SD 27.86 ± 7.18). Following artifact correction, previously rejected channels were reconstructed using spherical spline interpolation and the data were re-referenced to the common average reference. Finally, epochs were separated according to motor imagery condition (left- and right-hand imagery). A minimum of 90 artifact-free trials per condition was retained for each participant for subsequent analyses.

Following preprocessing, EEG data were analyzed using five complementary approaches. Time–frequency analysis was performed to verify that the motor imagery task successfully modulated sensorimotor oscillatory activity, as reflected by ERD. TEPs were analyzed to characterize cortical responses to stimulation. Source localization of group-averaged TEPs was conducted to visualize the spatiotemporal propagation of TEPs, and brain-state dependency analyses were performed to investigate the relationship between pre-stimulus oscillatory activity and TEP amplitudes.

First, time–frequency decomposition was performed using complex morlet wavelets applied to single-trial EEG data. Power was computed at 15 logarithmically spaced frequencies between 4 and 40 Hz, with the number of wavelet cycles increasing from 3 to 10 across frequencies. Power estimates were calculated separately for left- and right-hand motor imagery conditions. Power values were normalized to a pre-stimulus baseline period (−5000 to −4000 ms relative to motor imagery cue onset) and expressed as percentage change from baseline. This analysis was used to verify that the motor imagery task successfully induced sensorimotor ERD in the mu (8–12 Hz) and beta (13–30 Hz) frequency bands.

Second, TEPs were obtained by averaging preprocessed EEG epochs time-locked to the TMS pulse across the 62 scalp electrodes. TEP analyses focused on predefined latency windows derived from previous TMS–EEG studies (De Goede et al., 2020; Rogasch et al., 2017; Ilmoniemi & Kičić, 2010) and confirmed by inspection of the grand-average TEP waveform across all participants. To account for interindividual variability in peak latency, peak amplitudes were extracted within the following latency windows: N15:10–20 ms, P30: 25–35 ms, N45: 40–50 ms, P60: 55–65 ms.

Third, source localization was performed using Brainstorm (Tadel et al., 2011). Cortical sources were estimated using the default ICBM152 template anatomy. A forward head model was computed using Open MEEG boundary element modeling (BEM), and source activity was reconstructed using standardized low-resolution brain electromagnetic tomography (sLORETA). Noise covariance was estimated from the pre-stimulus baseline period (−5000 to −4000 ms). Source estimates were calculated from the grand-average TEPs to visualize the spatiotemporal propagation of cortical activity following TMS. Source maps were threshold for visualization purposes and displayed as relative source activity over the cortical surface. Source localization was performed as an additional visualization analysis and was not part of the preregistered statistical analyses.

Forth, brain-state dependency analyses were conducted to examine whether pre-stimulus sensorimotor oscillatory activity predicted variability in TEP amplitudes. Pre-stimulus mu-band (8–12 Hz) and beta-band (13–30 Hz) ERD were quantified over the left sensorimotor electrodes (C3, CP3, CP1), corresponding to the stimulated hemisphere, during the motor imagery period (−2000 to −10 ms relative to the TMS pulse). Relationships between pre-stimulus oscillatory activity and TEP peak amplitudes (N15, P30, N45, and P60) were assessed using Shepherd’s robust correlation. Bayesian Pearson correlations were additionally performed to quantify the strength of evidence for these associations.

Finally, an exploratory laterality index (LI) analysis was performed, trial-by-trial sensorimotor brain states were characterized using a laterality index (LI) derived from mu-beta band (8–30 Hz) ERD over homologous left (C3, CP3, CP1) and right (C4, CP4, CP2) sensorimotor electrodes during the motor imagery period (−2000 to −10 ms relative to the TMS pulse). The LI was calculated as:

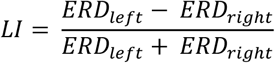

Negative LI values indicated greater ERD over the right sensorimotor cortex, whereas positive LI values indicated greater ERD over the left sensorimotor cortex in mu and beta band. For each participant, trials were classified according to the sign of the LI (negative vs. positive). This classification was independent of the instructed motor imagery condition and was based on the measured pre-stimulus sensorimotor activity. TEP peak amplitudes (N15, P30, N45, and P60) were averaged separately for negative and positive LI trials.

### Statistical Analysis

ERD differences between left- and right-hand motor imagery conditions were assessed using the cluster-based permutation test implemented in the FieldTrip toolbox (Oostenveld et al., 2011) with 2000 two-tailed permutations. Analyses were performed on ERD averaged across the left sensorimotor electrodes (C3, CP3, CP1). ERD was computed across frequencies ranging from 4 to 40 Hz and over the time interval from −2000 to −10 ms relative to the TMS pulse. Paired-samples t-tests were calculated at each time–frequency point to compare the two motor imagery conditions. Statistical significance was defined as a cluster-corrected p < .05.

To determine whether motor imagery influenced TEP amplitudes on the stimulated side (left M1), TEPs were averaged across the left sensorimotor electrodes (C3, CP3, CP1). Peak amplitudes were extracted within four latency windows (N15: 10–20 ms, P30: 25–35 ms, N45: 40–50 ms, and P60: 55–65 ms). Peak amplitudes were analyzed using a 2 × 4 repeated-measures ANOVA with Imagery Condition (left-vs. right-hand motor imagery) and TEP component (N15, P30, N45, and P60) as within-subject factors.

For brain-state dependency analyses, pre-stimulus sensorimotor activity was quantified as mu (8–12 Hz) and beta (13– 30 Hz) ERD over the left sensorimotor electrodes (C3, CP3, CP1) during the motor imagery period (−2000 to −10 ms relative to the TMS pulse). TEP peak amplitudes were extracted from the same electrodes within each predefined latency window. Relationships between pre-stimulus ERD and TEP amplitudes, as well as associations between TEP components, were assessed using Shepherd’s robust correlation (Schwarzkopf et al., 2012), which identifies bivariate outliers and computes correlation coefficients after their exclusion. Robust correlation coefficients (r) and corresponding p-values were reported.

To characterize trial-by-trial sensorimotor brain states, TEP peak amplitudes (N15, P30, N45, and P60) were compared between negative and positive laterality index (LI) trials using a two-way repeated-measures ANOVA with LI condition (negative vs. positive) and TEP component (N15, P30, N45, and P60) as within-subject factors.

To complement the frequentist analyses, bayesian statistics were performed in JASP (version 0.95.4). Bayesian paired-samples *t*-tests were conducted to compare mu- and beta-band ERD between left- and right-hand motor imagery conditions. Bayesian repeated-measures ANOVAs were performed for both the imagery condition × TEP component analysis and the exploratory LI condition × TEP component analysis. Bayesian Pearson correlations were additionally computed to quantify the evidence for associations between pre-stimulus oscillatory activity and TEP amplitudes. Bayes factors (BF_10_) are reported, with BF_10_ > 3 interpreted as moderate evidence for the alternative hypothesis and BF_10_ < 1/3 interpreted as moderate evidence for the null hypothesis.

## Results

Time–frequency analyses revealed pronounced event-related desynchronization (ERD) within both the mu (8–12 Hz) and beta (13–30 Hz) frequency bands during motor imagery (**Fig. *1***). Topographical maps averaged across the motor imagery period (−2000 to −10 ms before TMS) demonstrated bilateral sensorimotor ERD for both left- and right-hand motor imagery, with stronger desynchronization over the left hemisphere. The upper row of **Fig. *1***shows the scalp distribution of beta-band ERD, whereas the lower row depicts the corresponding mu-band ERD for both imagery conditions. Time–frequency representations averaged across the left sensorimotor electrodes (C3, CP3, CP1) revealed a transient increase in low-frequency power (4–8 Hz) following cue presentation, followed by sustained mu- and beta-band desynchronization throughout the motor imagery period until TMS delivery. Cluster-based permutation testing revealed no significant differences in oscillatory activity between left- and right-hand motor imagery over either the left (C3, CP3, CP1) or right (C4, CP4, CP2) sensorimotor electrode clusters (all cluster-corrected *p* > .05). To quantify the evidence supporting these null findings, Bayesian paired-samples *t*-tests were additionally performed. Moderate evidence favored the null hypothesis for both mu-band ERD (BF_10_ = 0.22, error% = 0.03) and beta-band ERD (BF_10_ = 0.20, error% = 0.03), indicating that the observed data were approximately 4.5 and 5.0 times more likely under the null hypothesis than under the alternative hypothesis, respectively. Together, the frequentist and Bayesian analyses indicate that left- and right-hand motor imagery produced comparable bilateral sensorimotor ERD and did not generate distinguishable oscillatory brain states. Consequently, data from both imagery conditions were pooled for the subsequent brain-state dependency analyses.

**Fig. 1:**
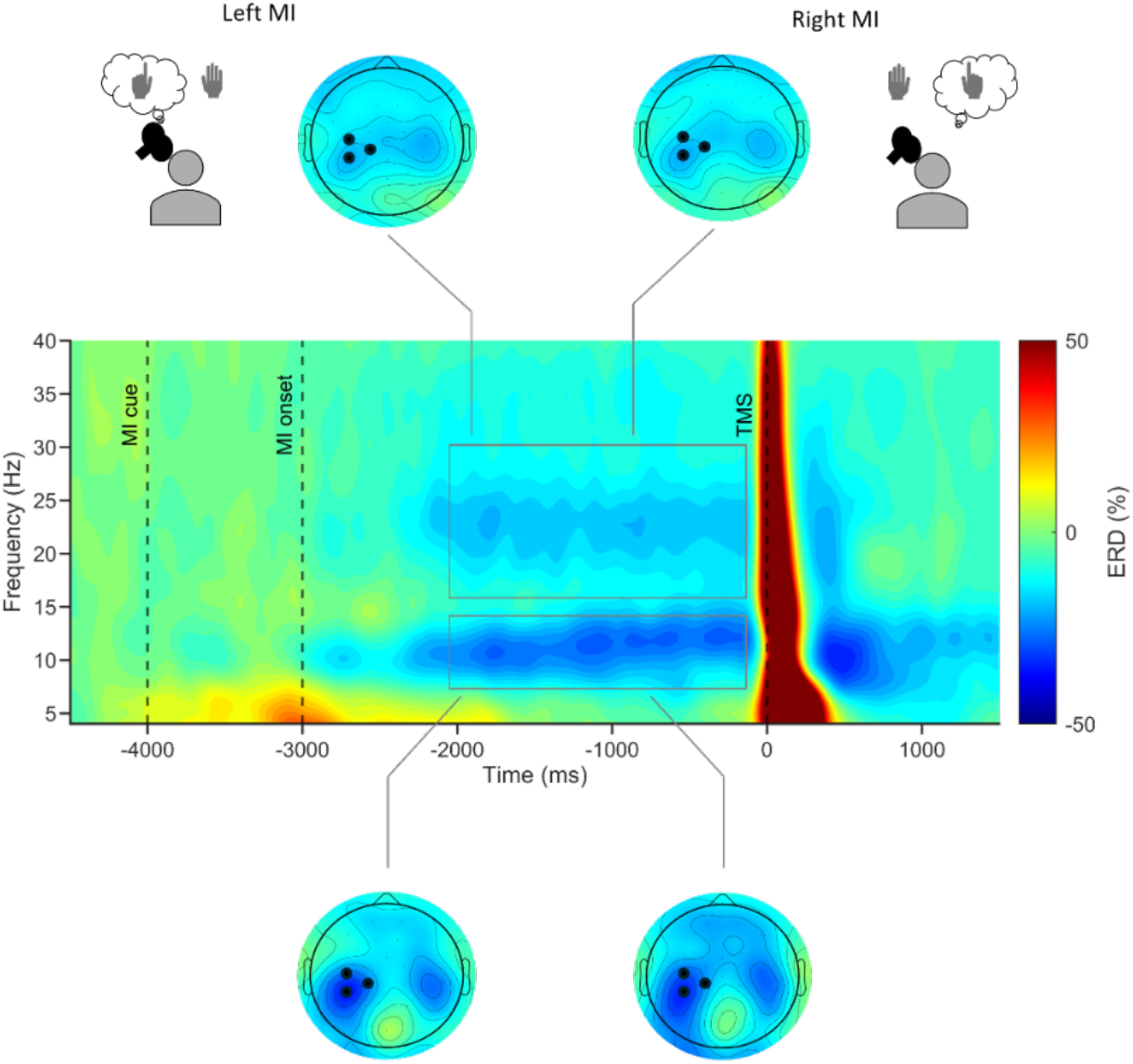
Event-related desynchronization during motor imagery. Topographical distribution of beta-band (13–30 Hz; top row) and mu-band (8–12 Hz; bottom row) ERD averaged across all participants (N = 28) during left-hand (left column) and right-hand (right column) motor imagery. ERD was averaged over the motor imagery period (−2000 to −10 ms relative to the TMS pulse; indicated by the black rectangle in the time–frequency plot). Both imagery conditions elicited bilateral sensorimotor desynchronization, with stronger ERD over the left hemisphere. The central panel shows the grand-average time–frequency pooled across left- and right-hand motor imagery conditions and averaged over the left sensorimotor electrodes (C3, CP3, CP1; indicated by black markers in the topographical maps). Dashed vertical lines indicate cue onset and motor imagery onset. Sustained ERD was observed within the mu and beta frequency bands throughout the motor imagery period. A transient increase in low-frequency power (4–8 Hz) was observed immediately following cue presentation. The same color scale is used for both the topographical maps and the time–frequency representation.

Fig. **2** illustrates the grand-average TEPs across all 62 EEG electrodes (bottom), scalp topographical distributions at representative latencies (middle), and source localization estimates of cortical activity propagation (top). Single-pulse TMS over the left primary motor cortex (M1) elicited clear and reproducible TEPs across participants. The butterfly plot revealed several prominent TEP components within the predefined latency windows of 10–20 ms, 25–35 ms, 40– 50 ms, and 55–65 ms. Early activity was maximal over the stimulated left sensorimotor cortex, whereas later components showed broader bilateral frontocentral distributions. Source reconstruction similarly indicated rapid propagation of TMS-evoked activity from the stimulated motor cortex to distributed bilateral motor-related networks. To determine whether motor imagery influenced TEP amplitudes, peak amplitudes of the N15, P30, N45, and P60 components were compared between left- and right-hand motor imagery conditions using a 2 × 4 repeated-measures ANOVA. A significant main effect of TEP Component was observed (*F*(3,81) = 9.55, *p* < .001), indicating that TEP amplitudes differed across latency windows. In contrast, there was no main effect of imagery condition (*F*(1,27) = 0.36, *p* = .55), indicating that overall TEP amplitudes were comparable between left- and right-hand motor imagery conditions. Although the component × condition interaction reached significance in the uncorrected analysis (*F*(3,81) = 2.98, *p* = .036), it was no longer significant after Greenhouse–Geisser correction (*p* = .056). Consistent with these findings, post hoc comparisons revealed no significant differences between imagery conditions for any TEP component (all *p* > .05). To quantify the evidence supporting the null findings, a bayesian repeated-measures ANOVA was additionally performed. The model including only the TEP component was the best-fitting model (BF_10_ = 1.00). Models additionally including imagery condition (BF_10_ = 0.329, error% = 4.60) or the imagery condition × TEP component interaction (BF_10_ = 0.346, error% = 4.08) received less support. These results indicate that the observed data were approximately 3 times more likely under models excluding imagery condition than under models including it, providing anecdotal to moderate evidence against an effect of imagery condition on TEP amplitudes.

**Fig. 2:**
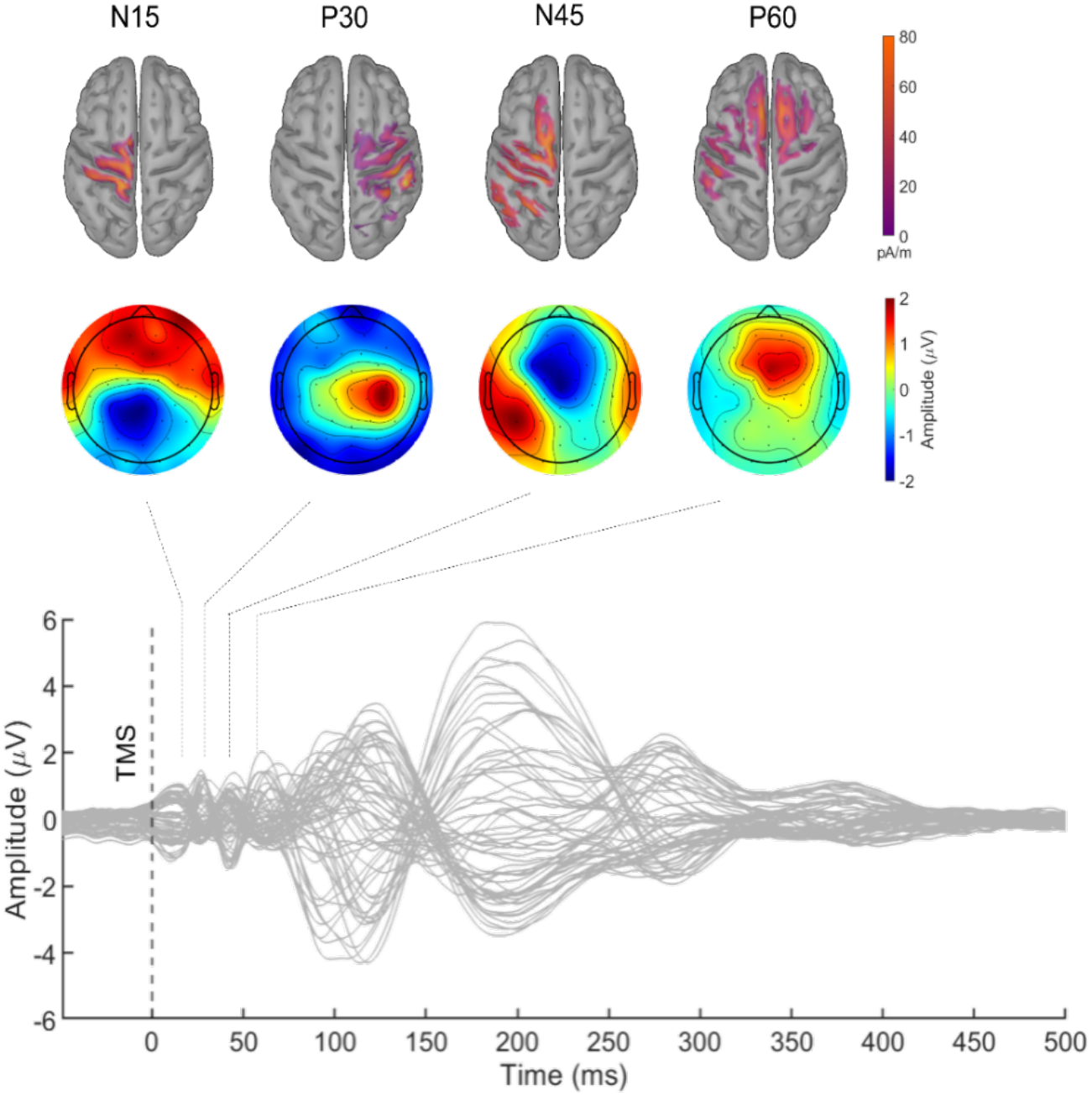
TEPs following stimulation of the left primary motor cortex. Grand-average TEPs pooled across left- and right-hand motor imagery conditions and averaged across all participants (N = 28). Bottom panel: Butterfly plot showing TEP waveforms from all 62 EEG channels. The vertical dashed line indicates TMS onset (0 ms). Middle panel: Scalp topographical distributions of TEP amplitudes post-stimulation latencies (10–65 ms), illustrating the spatiotemporal evolution of cortical activity following stimulation. Top panel: Source localization estimates reconstructed from the grand-average TEPs, demonstrating the propagation of TEPs from the stimulated left primary motor cortex, followed by activation of contralateral motor regions and subsequent recruitment of distributed bilateral premotor and motor-related cortical areas.

Correlation analyses performed on the left sensorimotor electrodes (C3, CP3, CP1) revealed a significant positive association between pre-stimulus beta-band ERD and the amplitude of the P60 component (r = 0.52, p = .015; Figure 3). No significant relationships were observed between pre-stimulus oscillatory activity and the earlier TEP components. In addition, pre-stimulus mu- and beta-band ERD were strongly correlated (r = 0.77, p < .001). Among the TEP components, a significant positive correlation was observed between the N45 and P60 components (r = 0.56, p = .009), whereas no other component pairs reached statistical significance. Bayesian correlation analyses yielded converging evidence for these findings. Decisive evidence supported the positive association between pre-stimulus beta-band ERD and P60 amplitude (BF_10_ = 197.39), indicating that the observed data were approximately 197 times more likely under the alternative hypothesis than under the null hypothesis. Strong evidence was observed for the correlation between mu- and beta-band ERD (BF_10_ = 26,316) and for the association between the N45 and P60 components (BF_10_ = 25,004). In contrast, the associations between beta-band ERD and the N15, P30, and N45 components all yielded BF_10_ values below 1, providing evidence in favor of the absence of these relationships.

**Fig. 3:**
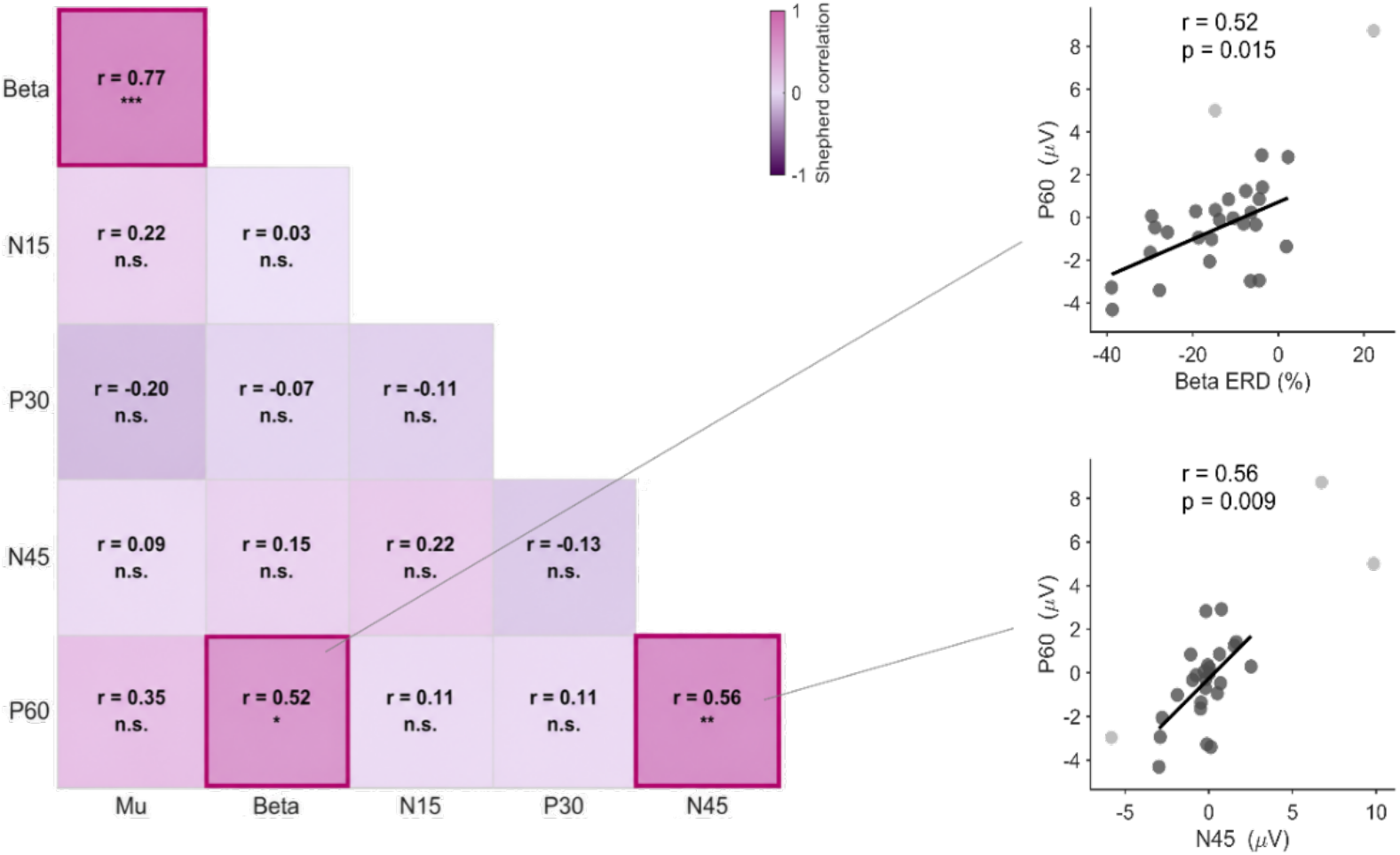
Relationship between pre-stimulus sensorimotor ERD and TEPs. Correlation analyses between sensorimotor ERD during the motor imagery period (−2000 to −10 ms relative to the TMS pulse) and TEP amplitudes obtained from the left sensorimotor electrodes (C3, CP3, CP1). Left: Shepherd’s robust correlation matrix showing pairwise associations among mu ERD (8–12 Hz), beta ERD (13–30 Hz), and TEP components (N15, P30, N45, and P60). Correlation coefficients (r) are displayed within each cell, and significant correlations are highlighted. Right: Scatter plots illustrating the association between pre-stimulus beta ERD and P60 amplitude and association between N45 and P60 amplitudes. Regression lines were fitted after exclusion of bivariate outliers according to Shepherd’s robust correlation procedure. Excluded bivariate outliers are shown as light gray data points.

**Fig. 4:**
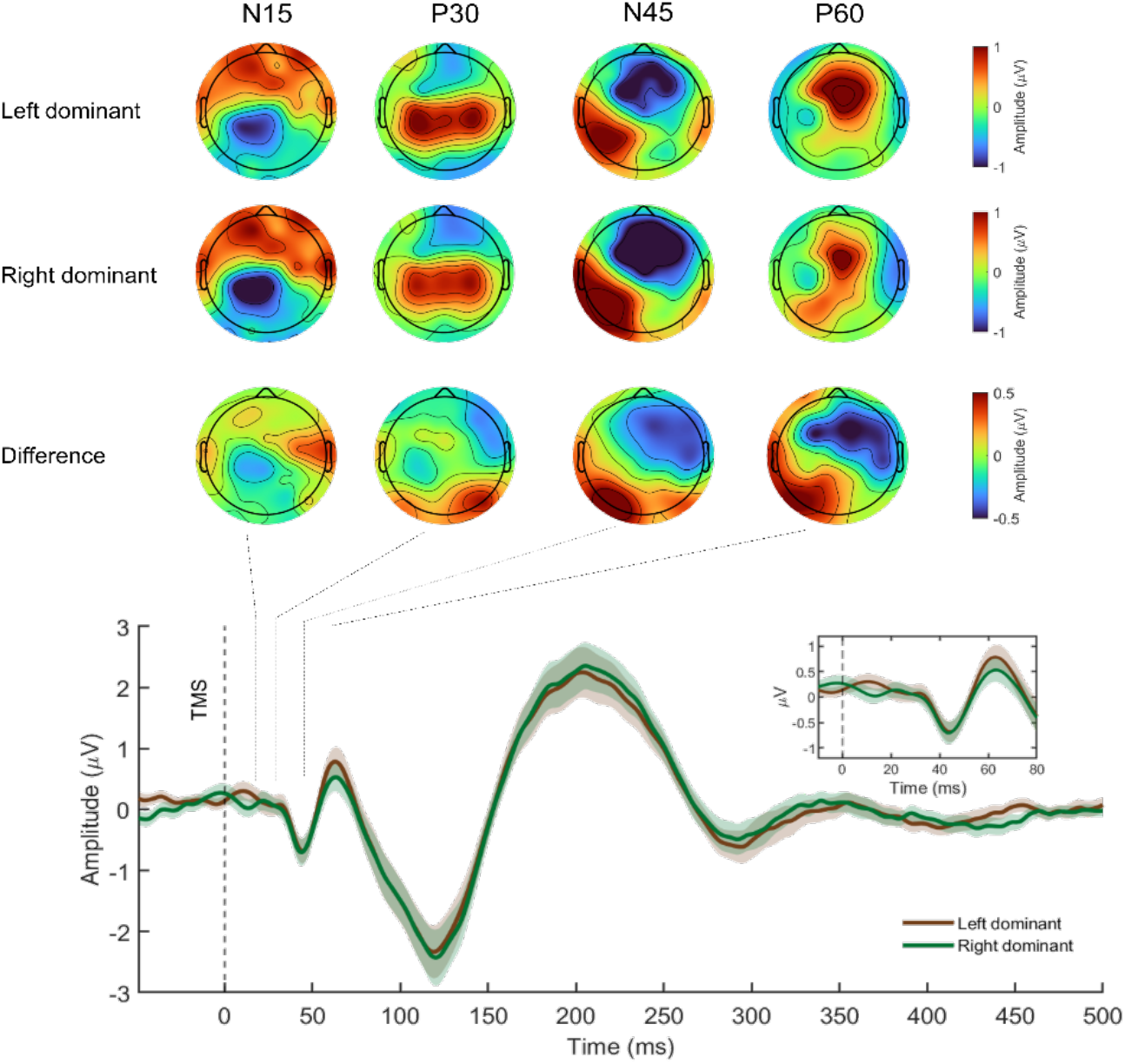
Trial-by-trial classification of TEPs based on sensorimotor laterality. Trials from both motor imagery conditions were pooled and subsequently classified according to a sensorimotor laterality index (LI), reflecting the relative asymmetry of mu-beta band activity between homologous left (C3, CP3, CP1) and right (C4, CP4, CP2) sensorimotor electrodes (*N* = 28). Trials were categorized as left-dominant or right-dominant based on the sign of the LI. Grand-average TEPs for left- and right-dominant trials, shown for the left sensorimotor electrodes and their right-hemisphere homologues. The vertical dashed line indicates the time of TMS onset (0 ms). Scalp topographies illustrating TEP amplitudes for left-dominant trials (top), right-dominant trials (middle), and their difference (bottom).

To investigate trial-by-trial variability in sensorimotor brain state, TEP amplitudes were compared between negative and positive laterality index (LI) trials using a 2 × 4 repeated-measures ANOVA with factors condition (negative vs. positive LI) and TEP component (N15, P30, N45, P60). A significant main effect of TEP component was observed (F(3,81) = 6.34, p = .001, Greenhouse–Geisser corrected), indicating differences in amplitude across latency windows. In addition, a significant main effect of LI condition was found (F(1,27) = 5.04, p = .033), suggesting overall differences in TEP amplitudes between negative and positive LI trials. However, the component × condition interaction was not significant (F(3,81) = 0.87, p = .41, Greenhouse–Geisser corrected), indicating that the effect of LI condition did not differ reliably across TEP components. To further explore potential component-specific effects, post hoc paired-samples t-tests were conducted and corrected for multiple comparisons using Bonferroni correction. These analyses revealed that only the P60 component remained significantly modulated by LI condition (p = .046), whereas no significant differences were observed for N15, P30, or N45 (all corrected p ≥ .135).

## Discussion

The present study investigated whether mentally induced sensorimotor brain states modulate cortical responses to TMS. Although motor imagery successfully induced robust mu- and beta-band ERD over sensorimotor regions, left- and right-hand motor imagery did not produce distinguishable patterns of oscillatory activity. This indicates that the two imagery conditions elicited comparable sensorimotor brain states at the stimulated cortex. Consistent with this, TEP amplitudes did not differ between conditions. However, pre-stimulus sensorimotor oscillatory activity was associated with variability in a later TEP component (P60), suggesting that ongoing brain activity may selectively influence later stages of TMS-evoked cortical processing.

Although the imagery task involved unilateral index finger tapping, ERD was observed bilaterally over sensorimotor regions. This finding is consistent with previous motor imagery studies demonstrating that unilateral motor imagery engages distributed sensorimotor networks in both hemispheres rather than producing strictly lateralized cortical activation (Hasegawa et al., 2017; Pfurtscheller et al., 2006; Pfurtscheller & Neuper, 1997; Zich, Harty, et al., 2017). While ERD is typically stronger over the hemisphere contralateral to the imagined movement, ipsilateral sensorimotor areas are also recruited during motor imagery, reflecting the involvement of premotor, supplementary motor, and interhemispheric motor networks. Consistent with this interpretation, Beck et al., (2025) reported bilateral sensorimotor desynchronization during unilateral voluntary movements, suggesting that both overt movement and motor imagery involve widespread modulation of motor cortical networks. The bilateral ERD observed in the present study therefore suggests that both imagery conditions induced broadly similar sensorimotor states at the stimulated left motor cortex, which may explain the absence of condition-specific differences in TEP amplitudes.

The most important finding of the present study was that pre-stimulus beta-band ERD was associated with the amplitude of the P60 component. Specifically, lower pre-stimulus beta ERD was associated with larger P60 amplitudes. This association was observed across participants and therefore reflects an individual-differences relationship rather than direct evidence for trial-to-trial modulation of TEPs. Nevertheless, the finding suggests that differences in the pre-stimulus sensorimotor state are related to variability in later TEP. Interestingly, the association was restricted to the P60 component, whereas earlier TEP components were not significantly related to pre-stimulus oscillatory activity. This pattern raises the possibility that later stages of TEP may be more sensitive to differences in the functional state of the motor system than the initial cortical response. Such an interpretation is consistent with previous TMS–EEG studies showing that later TEP components increasingly reflect recurrent cortico-cortical interactions and large-scale network processing rather than the direct local response to stimulation (De Goede et al., 2020; Ilmoniemi et al., 1997; Ilmoniemi & Kičić, 2010; Komssi et al., 2002). To further account for trial-by-trial variability in sensorimotor brain state, we performed an exploratory analysis in which trials were classified based on a sensorimotor laterality index (LI), reflecting the relative balance of activity between the two hemispheres. This analysis was motivated by the observation that the instructed motor imagery conditions did not produce clearly distinguishable brain states at the cortical level. Consistent with the correlation results, LI-based trial classification revealed that differences in TEP amplitudes were primarily observed for later components, with the most robust effect for the P60 component. In contrast, early TEP components were not reliably modulated by LI. Together, these findings suggest that, trial-by-trial fluctuations in sensorimotor brain state rather than the instructed motor imagery condition are associated with variability in later stages of TMS-evoked cortical processing. Notably, the absence of a significant interaction in the ANOVA suggests that these component-specific effects should be interpreted with caution. However, the convergence between the LI-based analysis and the correlation results strengthens the conclusion that later TEP components, particularly P60, are sensitive to ongoing brain-state fluctuations.

Notably, only beta ERD was associated with P60 amplitude, whereas no significant relationship was observed for mu ERD. Previous MEG studies suggest that mu and beta rhythms originate from partially distinct sensorimotor generators, with beta activity showing stronger contributions from precentral motor regions and mu activity being relatively more prominent in postcentral somatosensory regions (Cheyne, 2013). Because TMS was delivered over the left primary motor cortex, beta-band activity may have provided a more direct measure of the functional state of the stimulated cortical region. In addition, beta oscillations have been strongly linked to motor cortical excitability and the maintenance of the current motor state, further supporting their relevance for predicting TEPs. The absence of a significant mu–TEP relationship therefore suggests that state-dependent modulation of TEPs may be more closely related to motor cortical beta dynamics than to sensorimotor mu activity (Ahola et al., 2025; Sarasquete et al., 2026). Importantly, the beta–P60 relationship was observed only in the stimulated hemisphere. The same analyses were performed for the homologous right sensorimotor electrodes (C4, CP4, CP2). No significant associations were found between pre-stimulus ERD and any TEP component (all p > .05). The only significant correlation observed was between pre-stimulus mu and beta ERD (r = 0.45, p = .046). This hemispheric specificity suggests that the observed effect was not driven by global fluctuations in oscillatory activity but rather reflected the local functional state of the cortical region directly perturbed by TMS.

Finally, the positive association between N45 and P60 amplitudes suggests that successive stages of TMS-evoked processing may not be independent but rather reflect the propagation of activity through interconnected motor networks. This interpretation is supported by the source localization results, which indicated an initial activation of the stimulated motor cortex followed by rapid propagation toward contralateral and bilateral motor-related cortical regions. Similar spatiotemporal propagation patterns have been reported previously and are thought to reflect the recruitment of distributed cortico-cortical motor networks following TMS (Beck et al., 2025; Ilmoniemi & Kičić, 2010).

## Limitations

Several limitations should be considered when interpreting the present findings. First, the study was conducted in a relatively small sample of healthy young adults, which may limit the generalizability of the results to other age groups and clinical populations. Second, TMS was applied only over the left primary motor cortex at a single stimulation intensity (80% RMT). It therefore remains unclear whether similar brain-state-dependent effects would be observed at different stimulation intensities or at other cortical targets. Third, although motor imagery successfully induced robust sensorimotor ERD, desynchronization was observed bilaterally, resulting in largely similar oscillatory states across the two imagery conditions. Future studies combining real-time monitoring of oscillatory activity with neurofeedback or brain-state-dependent stimulation approaches may help achieve more consistent modulation of sensorimotor brain states and further clarify their influence on TEPs. Although the exploratory laterality analysis incorporated trial-level variability, the present study did not directly model continuous within-subject trial-to-trial relationships between pre-stimulus oscillatory activity and TEP amplitude. Future studies using single-trial TEP estimation and hierarchical modelling could more directly test this form of brain-state dependency.

## Conclusion

In summary, motor imagery successfully induced robust mu- and beta-band desynchronization over sensorimotor regions, confirming effective modulation of sensorimotor brain state. Consistent with previous motor imagery studies, desynchronization was observed bilaterally rather than being strictly lateralized to the imagined movement. Despite these pronounced oscillatory changes, motor imagery did not significantly alter TMS-evoked potentials, suggesting that early cortical responses to TMS remain relatively stable across behaviorally induced sensorimotor states. Pre-stimulus beta-band ERD was significantly associated with the amplitude of the later P60 component, indicating that ongoing sensorimotor brain activity may influence subsequent stages of TMS-evoked network processing. The absence of similar relationships for earlier TEP components suggests that state-dependent effects may emerge primarily during later cortico-cortical processing rather than during the initial local response to stimulation. Together, these findings highlight the importance of ongoing brain state in shaping TMS-evoked cortical responses and support the use of TMS–EEG as a tool for investigating brain-state-dependent dynamics of the human motor system. Future studies should examine whether these relationships are influenced by stimulation intensity and whether similar state-dependent effects are present in clinical populations, where pre-stimulus oscillatory activity may provide useful information for predicting responsiveness to TMS interventions.

## Author Contributions

MD designed the study, collected the data, implemented the analysis scripts, performed the analyses, interpreted the results, and wrote the main manuscript text. SD contributed to the study design, supervised the project, and revised the manuscript. All authors reviewed the manuscript, and approved the final submitted version.

## Ethics Statement

The study was reviewed and approved by the ethics committee of the *Carl von Ossietzky University of Oldenburg* (Drs. No. EK/2024/057), Oldenburg Germany. The participants provided their written informed consent to participate in this study.

## Data Availability Statement

The raw datasets used for the current study are available by contacting Motahare Delbari on request.

## Statements and Declarations

CSH holds a patent on transcranial brain stimulation. All other authors declare no conflicts of interest.

## Acknowledgements

We thank Reiner Emkes for technical support and Carlotta Wall for assistance with data collection. We also thank all participants for their time and commitment to this study.

